# Spinal motoneuron firing properties mature from rostral to caudal during post-natal development of the mouse

**DOI:** 10.1101/2020.05.15.097436

**Authors:** Calvin C. Smith, Robert M. Brownstone

## Abstract

Altricial mammals are born with immature nervous systems comprised of circuits that do not yet have the neuronal properties and connectivity required to produce future behaviours. During the critical period of post-natal development, neuronal properties are tuned to participate in functional circuits. In rodents, cervical motoneurons are born prior to lumbar motoneurons, and spinal cord development follows a sequential rostro-caudal sequence. Here we asked whether birth order is reflected in the post-natal development of electrophysiological properties. We show that motoneurons of both segments have similar properties at birth and follow the same developmental profile, with maximal firing increasing and excitability decreasing into the 3^rd^ post-natal week. However, these maturative processes occur in cervical prior to lumbar motoneurons, correlating to the timing of arrival of descending systems. These results suggest that motoneuron properties do not mature by cell autonomous mechanisms alone, but rather depend on developing descending and spinal circuits.

## Introduction

Altricial mammals are born with immature nervous systems comprised of circuits that have neither the neuronal properties nor connectivity required to produce future behaviours. During the critical period of post-natal development, neuronal properties are tuned to participate in functional circuits. The degree to which these mature electrophysiological properties are determined by cell-intrinsic vs extrinsic (e.g. circuit) factors is not clear. To understand how neural circuits are ultimately tuned requires understanding of how properties of their component neurons develop.

A model system in which to understand this is the spinal cord, where two distinct regions – the cervical and the lumbar cord – include homologous circuits, with both regions containing circuitry to coordinate intra- and inter-limb movements required for locomotion (Jessell, 2000a; Goulding, 2009). The output from each region is mediated by motoneurons, which have overlapping molecular profiles in the two regions. During embryonic development, cervical motoneurons are born a few days prior to lumbar motoneurons in mice, rats, and chicks (Nornes and Das, 1974; Hollyday and Hamburger 1977), and these regions develop sequentially from rostral to caudal (Sagner & Briscoe, 2019). Is this developmental trajectory maintained in the post-natal critical period such that electrophysiological properties of cervical motoneurons are more mature at birth and then fully mature prior to lumbar motoneurons? Or do the development and maturation of these properties rely on the development of circuits, such as descending and sensory inputs, in which case the two populations would be born with similar properties that mature as movement develops during the post-natal critical period?

There is little question that both behaviour and motoneuron properties develop post-natally. Rodents are essentially immobile in the first 2 days following birth, until forelimb propelled pivoting and crawling become the dominant forms of ambulation during the first week of life. (Altman & Sudarshan, 1975; Sechzer *et al.*, 1984). Quadrupedal locomotion subsequently emerges around P10-12 when hindlimbs consistently support the weight of the lower quadrant (Jiang et al 1999), with locomotor maturity reached by the end of the 3^rd^ post-natal week (Altman & Sudarshan, 1975). And lumbar motoneuron properties change at least in the first post-natal week, a period (and region) in which electrophysiological experiments are usually done (Fulton & Walton, 1986; Nakanishi & Whelan, 2010; Quinlan *et al.*, 2011). But do these changes in properties proceed from rostral to caudal?

Many aspects of spinal circuit development ensue from rostral to caudal. Descending supraspinal pathways arrive and mature in cervical segments earlier than in lumbar segments. In rodents, fibres originating in the brainstem are the first to arrive in the spinal cord, sprouting into the grey matter of the cervical cord prenatally, and reaching the lumbar spinal segments at or shortly after birth (Vinay *et al.*, 2000). Descending modulatory systems follow a similar pattern: serotonergic innervation of the cervical spinal cord displays the adult profile by P14, whereas the fibre density in the lumbar cord does not mature until P21 (Bregman, 1987). Similarly, corticofugal axons innervate the cervical grey matter at P5-6 and arrive at the lumbar cord at approximately P9. But it is not until after P21 that all segments display the mature density and pattern of descending innervation (Donatelle, 1977; Gianino *et al.*, 1999). Sensory innervation of the spinal cord follows a similar trend, as cutaneous reflexes can be evoked in the muscles of the neck and forelimbs (E16-17) prior to in hindlimbs (E17-18) (Narayanan *et al.*, 1971). Furthermore, postnatal refinement of proprioceptive afferent input to motoneurons is dependent upon the maturation of descending input in the spinal cord and does not mature until the end of the 3^rd^ postnatal week in rats (Smith *et al.*, 2017). Thus, key spinal circuit development occurs in the first 3 post-natal weeks and, where studied, proceeds from the cervical to the lumbar spinal cord.

Do motoneuron properties follow this same pattern? In addition to being born earlier, anatomical data indicate that cervical motoneurons reach adult motoneuron size prior to lumbar motoneurons (Cameron *et al.*, 1989). In rats, the appearance of spontaneous burst activity in embryonic cervical motoneurons precedes that in lumbar motoneurons by about one embryonic day (Nakayama *et al.*, 1999). Importantly, it is clear that the transcriptional profile rather than circuit milieu (and limb movement) is sufficient for basic motoneuron properties to develop: motoneurons derived in culture from embryonic stem cells develop electrophysiological properties characteristic of spinal motoneurons (Miles et al., 2004).

We thus reasoned that, not only would motoneuron properties develop in the early post-natal period, but that if transcriptional profile alone were responsible for this maturation, then cervical motoneurons would be more mature at the time of birth and reach adult-like properties earlier than lumbar motoneurons. On the other hand, if circuit properties and limb movement are critical for maturation, then cervical and lumbar motoneurons would be born with similar properties that mature post-natally. To study this question, we used whole cell patch clamp recordings of mouse cervical and lumbar motoneurons to define and compare their electrophysiological properties at three time points during this critical period of development (P2-3, P6-7 & P14-21). We demonstrate that the two populations are born with similar properties and then develop through the third post-natal week. Furthermore, cervical motoneuron properties mature earlier than those of lumbar motoneurons. We thus suggest that transcriptional profiles alone are insufficient for this maturation, and that the post-natal development of circuits contributes to the tuning of neuronal properties.

## Results

To effect movement, motoneurons must have the machinery to integrate synaptic inputs and convert them to trains of action potentials of appropriate frequencies for the intended behaviour. The ability to do this processing arises from a combination of passive, transition, and repetitive firing properties. Here, we report our findings on post-natal development in each of these categories, comparing limb-innervating motoneurons within the cervical and lumbar spinal cord. For all analyses, we first assessed if there was an effect of age on the development of these properties in cervical and in lumbar motoneurons separately. Unless there was no effect of age on either segment, we next compared the 6 groups with each other (2 regions, each at 3 ages) using corrected pairwise comparisons.

### Postnatal development of motoneuron passive properties

The key passive properties to consider are resting membrane potential, whole cell capacitance, and input resistance. The resting membrane potential did not change between ages for either segment (Figure 1A, A’), suggesting that the potassium equilibrium potential and permeability of channels open at this potential are mature at birth (Lumbar: χ^2^ = 5.3, p= 0.070; Cervical: 1 way ANOVA, F=0.001, p=0.97).

**Figure 1.**
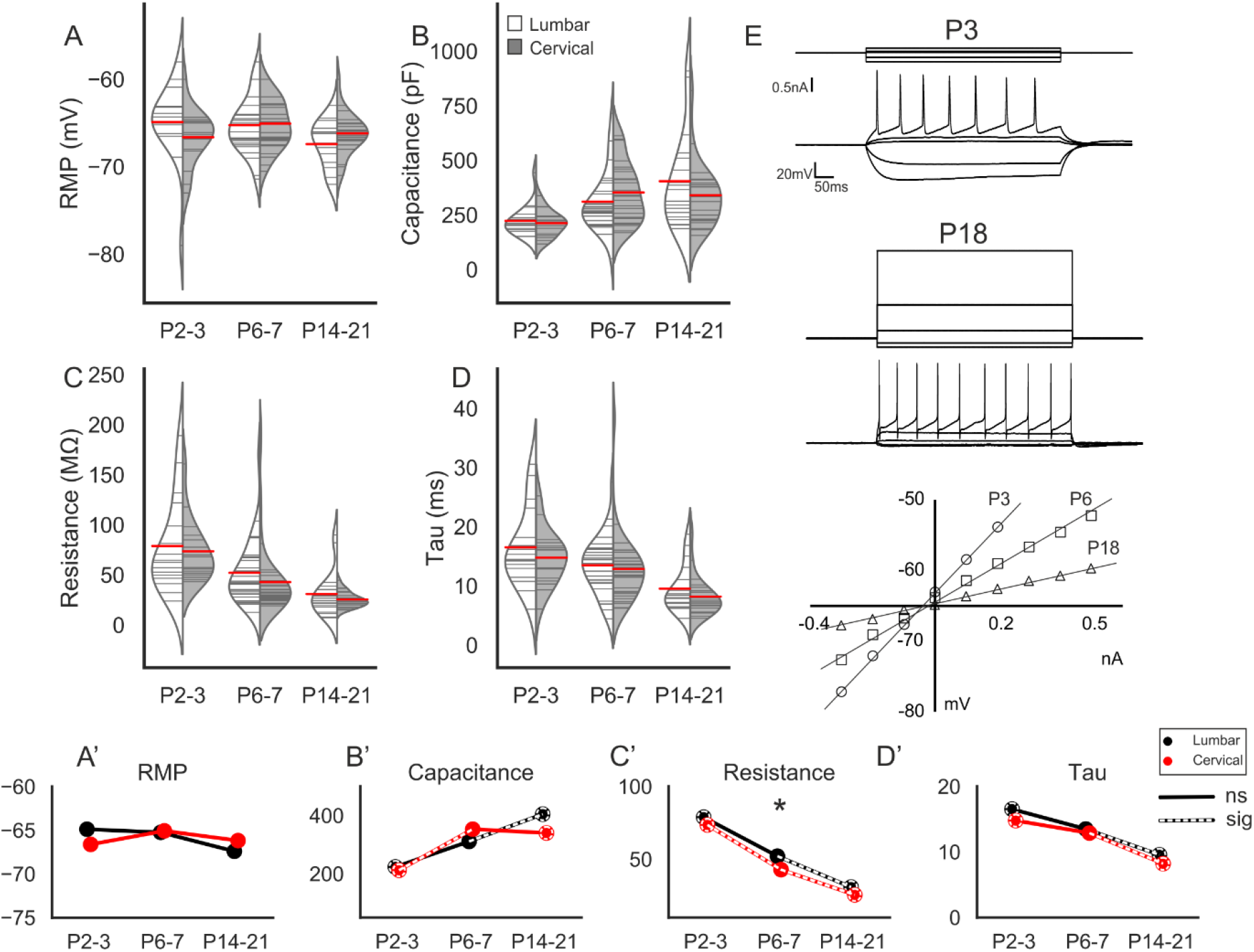
Postnatal development of motoneuron passive properties. (A-D) Split violin plots showing development of passive membrane properties. Grey horizontal lines within violins show individual observations for each age and location (cervical-white and lumbar-grey). The red horizontal lines represent mean values. (E) Membrane voltage responses to depolarising current pulse from −0.3nA in a P3 (Top) and P18 (middle) motoneuron. For both cells the top trace is the injected current and the bottom trace the membrane voltage response. Lower: A graph showing peak membrane voltage response (y-axis) to current input (x-axis) for a representative motoneuron within each age group. (A’-D’) Joint-plots of mean values illustrating the developmental profile of each measure in A-D. Over-laid white dashed lines represent statistically significant changes between age groups. Dashed circles represent statistically significant difference between P2-3 and P14-21. * represents a statistically significant difference between segments for a particular age group.

Whole cell capacitance (WCC) reflects neuronal size, which for motoneurons has implications on behavioural output, including muscle fibre types innervated and recruitment order. In lumbar motoneurons, there was a significant effect of age on development (KW χ^2^= 12.7, p= 0.001, Figure 1B, B’). WCC remained stable through the first post-natal week (P2-3=223 +/− 71 pF vs P6-7=311 +/− 140 pF, p=0.101), with the increase being thereafter (P14-21= 404 +/− 217 pF, p=0.048). The increase in WCC in cervical motoneurons was evident earlier, between P2-3 (212 +/− 59 pF) and P6-7 (353 +/− 136 pF, p=0.001), and remained stable afterwards (P14-21, 340 +/− 148 pF, p=0.456). There was no difference in membrane capacitance between cervical and lumbar motoneurons at any age (P2-3, p=0.917; P6-7, p=0.071 and P14-21, p=0.419). Together, these data suggest that both cervical and lumbar motoneurons increase in size post-natally, with the increase being evident earlier in cervical motoneurons.

Measurements of input resistance reflect the conductance properties of channels that open near resting potential, and also correlate with neuronal size. There was a significant developmental decrease in resistance in both lumbar (KW χ^2^= 17.2, p=1.8e-03) and cervical motoneurons (KW χ^2^= 35.7, p=1.8e-08, Figure 1C, C’, E). In lumbar motoneurons, there were no differences in resistance between P2-3 and P6-7 (P2-3=79 +/− 47 MΩ, vs P6-7=52 +/−32 MΩ, p=0.311), with the decrease occurring later, P6-7 vs P14-21=31 +/− 23 MΩ, p=5.3e-03). In cervical motoneurons there was an initial decrease in the first week that continued subsequently (P2-3, 74 +/− 31 MΩ vs P6-7, 43 +/− 34 MΩ, p=4.4e-07 and P6-7 vs P14-21, 26 +/− 8 MΩ, p=0.010). Comparing the resistances of cervical and lumbar motoneurons for each age group showed that they had similar start and end points, but differed at the middle time point. That is, there was no difference between regions at P2-3 (p=0.918) or P14-21 (0.918), but cervical motoneurons had a lower mean resistance at P6-7 (p=6.8e-05). This indicates post-natal maturation of these conductances in both groups, with cervical motoneurons maturing earlier.

Neuronal time constant, τ, combines WCC and input resistance, and reflects integrative properties of the cell. Analysis of τ in lumbar and cervical motoneurons revealed a developmental pattern similar to that seen for input resistance, with both decreasing significantly over time (Lumbar: KW χ^2^= 13.5, p= 0.001; Cervical: KW χ^2^= 22.7, p= 1.1e-05, Figure 1D, D’). τ did not change between P2-3 (16 +/− 6 ms) and P6-7 (13 +/− 4 ms, p=0.246) for lumbar motoneurons, but decreased between subsequent age groups (P6-7 vs P14-21= 9 +/− 3 ms, p= 0.008). Cervical motoneurons followed the same profile as τ decreased between P6-7 and P14-21 (P6-7= 12 +/− 5 ms vs P14-21= 8 +/− 3 ms, p=0.003), but not P2-3 (14 +/− 4 ms) and P6-7 (8 +/− 3 ms, p=0.096) (Figure 1 D, D’). There was no difference between cervical and lumbar motoneurons at any age (P2-3, p= 0.683; P6-7, p=0.370; P14-21, p=0.243). Together, these data indicate that the passive properties of motoneurons are similar by the third post-natal week, but cervical motoneurons reach this point prior to lumbar motoneurons.

### Postnatal development of motoneuron transition properties: action potentials

A key indicator of functional maturation is action potential (and after potential) morphology. The fast spike can be characterised by the ½ width, rate of rise and fall, threshold, amplitude, and depth of fall or fAHP. To assess changes in these characteristics we evoked single APs using 20ms current pulses while the resting membrane potential was held at −65mV. For both lumbar and cervical motoneurons, there was a significant decrease in action potential ½ width with age (Lumbar: ANOVA, F=39.52, p= 2.8e-08, Cervical: KW χ^2^= 32.59, p= 8.345e-08, Figure 2A, A’).

**Figure 2.**
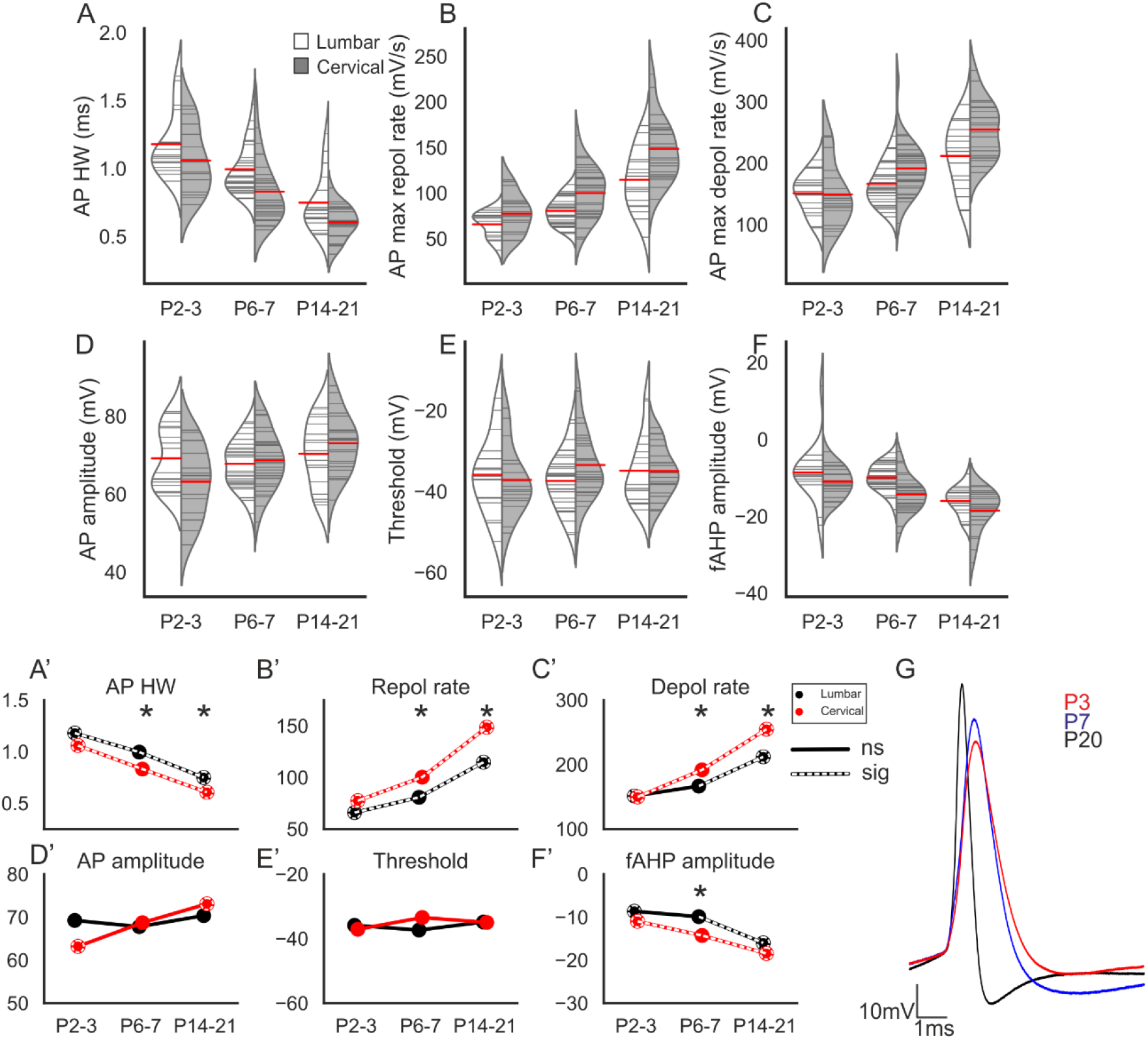
Postnatal development of action potential characteristics. (A-F) Split violin plots illustrating the changes in action potential characteristics at each age in lumbar (white) and cervical (grey) motoneurons. (A’-F’) Joint-plots of means illustrating the developmental profile of each measure. Over-laid white dashed lines represent statistically significant changes between age groups. Dashed circles represent statistically significant difference between P2-3 and PI4-21. ^*^ represents a statistically significant difference between segments for a particular age group. (G) Overlaid traces from representative Mns from each of the age groups. Each tracc was averaged from 15-30 action potentials with the cells resting membrane potential held at −65 mV.

In lumbar motoneurons, the AP half width reduced between each time point (P2-3=1.2 +/− 0.2 ms vs P6-7=1.0 +/− 0.2 ms, p=0.003; P6-7 vs P14-21= 0.7 +/− 0.2ms, p=3e-03). The decrease in half width between P2-3 and P6-7 was associated with an increase in the rate of repolarisation (max repolarisation rate: P2-3=66 +/− 14 mV/s vs P6-7= 81 +/− 15 mV/s, p=0.007, Figure 2B, B’) – there was no difference in the maximum depolarisation rates between these age groups (P2-3=152 +/− 33, P6-7=166 +/− 34, p=0.287, Figure 2C, C’). Between P6-7 and P14-21, however, there was an increase in both the maximum rates of depolarisation (P14-21=212 +/− 53 mV/s, p=0.003) and repolarisation (P14-21=114 +/− 34 mV/s, p=8e-3). These data suggest that delayed rectifier potassium channels mature earlier than voltage-gated sodium channels.

Action potential half width also decreased in cervical motoneurons between P2-3 (1.1 +/− 0.2ms) and P6-7 (0.8 +/− 0.2ms, p=0.001) and between P6-7 and P14-21 (0.6 +/− 0.1, p=0.001). Unlike lumbar motoneurons, both maximum rate of depolarisation (P2-3=150 +/− 51 mV/s vs P6-7=192 +/− 44 mV/s, p= 0.003; P6-7 vs P14-21 255 +/− 46 mV/s, p= 3.3e-06) and repolarisation (P2-3=77 +/− 22 mV/s vs P6-7=100 +/− 28 mV/s, p= 0.008; P6-7 vs P14-21 149 +/− 34.7mV/s, p= 2.4e-05) increased through this period and thus contributed to the reduction in AP half width between each age group. These data suggest that both delayed rectifier and voltage-gated sodium channels mature throughout this period in cervical motoneurons.

At P2-3 no differences between cervical and lumbar motoneurons were observed in action potential half width (p=0.151), or maximum rate of depolarisation (p= 0.918) or repolarisation (p=0.140). At P6-7, cervical motoneurons had shorter duration action potentials compared to lumbar motoneurons due to higher rates of both depolarisation and repolarisation (action potential half width: p=0.001, maximum rate of depolarisation: p= 0.036 and maximum rate of repolarisation: p=0.005). This was also true of cervical motoneurons in the P14-21 group (action potential half width p=0.010, maximum rate of depolarisation: p=4e-03, maximum rate of repolarisation: p=0.003), again suggesting an earlier maturation of cervical motoneurons.

We next assessed changes in action potential amplitude by measuring the difference between peak voltage and voltage threshold. There was no effect of age on AP amplitude for lumbar motoneurons (ANOVA, F=0.03, P=0.863), however there was a significant increase for cervical motoneurons (ANOVA, F=13.95, p= 3.93e-03, Figure 2 D, D’).

The increase in AP amplitude in the cervical cord was gradual as there was no difference between P2-3 (63 +/− 9 mV) and P6-7 (69 +/− 7.4 mV, p= 0.070) or P6-7 and P14-21 (73 +/− 7.7 mV, p= 0.109), however P14-21 was significantly greater than P2-3 (p=0.001). These data suggest that action potential conductances mature throughout this period in cervical motoneurons.

Because spike amplitude is measured as the difference between threshold and peak, threshold changes could contribute to changes in amplitude. However, we found no effect of age on spike threshold for either lumbar (F=0.689, p=0.410) or cervical motoneurons (F= 0.206, p=0.652, Figure 2E, E’), suggesting that changes are due to overshoot (Figure 2G).

Interestingly, though, there were no differences in AP amplitude between cervical and lumbar motoneurons at any age (Lumbar: P2-3 = 69 +/− 7.8 mV, p= 0.070; P6-7 = 68 +/− 6.1 mV. p=0.800; P14-21 = 70.4 +/− 8.5 mV, p=0.173). Although there is a gradual increase in amplitude with age for cervical motoneurons, the apparent lack of developmental change for lumbar motoneurons and lack of difference between segments suggests that development of Na^+^ may not contribute as much, relative to later activated channels, to the development of spike morphology.

The amplitude of the fAHP (measured from threshold to trough), which largely reflects conductances of voltage-gated potassium channels involved in repolarisation, increased with postnatal development for both lumbar (KW χ^2^=26.11, p= 2.13e-06) and cervical (ANOVA, F=23.48, p= 7.81e-06, Figure 2F, F’, G) motoneurons. In lumbar motoneurons, there was no difference in amplitude between P2-3 (−8.6 +/− 7.5 mV) and P6-7 (−9.9 +/− 3.5 mV, p= 0.410), but it increased between P6-7 and P14-21(−16 +/− 3.6 mV, p=2.0e-06). In cervical motoneurons, fAHP increased between P2-3 (−11 +/− 4.1 mV) and P6-7 (−14 +/− 4.8 mV, p= 0.022), and between P6-7 and P14-21 (19 +/− 5.5 mV, p= 0.022). Again, the only difference between spinal segments was a significantly greater fAHP in cervical motoneurons compared to lumbar motoneurons at P6-7 (P6-7, p=2.9e-03; P2-3, p= 0.272; P14-21, p= 0.162). This is further evidence that motoneuron voltage-gated potassium channels involved in repolarisation develop significantly post-natally in both segments, and that the process occurs earlier in cervical motoneurons.

### Postnatal development of motoneuron transition properties: after potentials

Action potential after potential characteristics contribute to the repetitive firing properties of motoneurons (Granit *et al.*, 1963a; Kernell, 1979). The slower after potentials such as the medium after hyperpolarisation (mAHP) and after depolarisation (ADP) contribute significantly to repetitive firing capabilities of MNs, and reflect motoneuron type (Granit *et al.*, 1963b; Kernell, 1972; Kernell & Monster, 1982). For both lumbar and cervical motoneurons, mAHP amplitude (measured from RMP to trough, Figure 3A, A’) was significantly reduced with development (Lumbar: KW χ^2^=13.6, p=0.001; Cervical: KW χ^2^= 11.6, p= 0.002). In lumbar motoneurons, there was no difference between P2-3 (5.3 +/− 2.4mV) and P6-7 (4.1 +/− 1.9mV, p=0.122) or P6-7 and P14-21 (3.0 +/− 1.6mV, p=0.093), but there was a reduction between P2-3 and P14-21 (p=0.005), suggesting a gradual reduction in amplitude over time. The same pattern of development was observed in cervical motoneurons (P2-3=5.4 +/− 1.9 mV vs P6-7=4.4 +/− 2.6 mV, p=0.128; P6-7 vs P14-21=3.3 +/− 1.3 mV,p=0.169; P2-3 vs P14-21, p=0.004). Moreover, there were no differences between cervical and lumbar motoneuron mAHPs at any of the age groups (P2-3, p=0.810; P6-7, p=0.942; P14-21, p=0.527).

**Figure 3.**
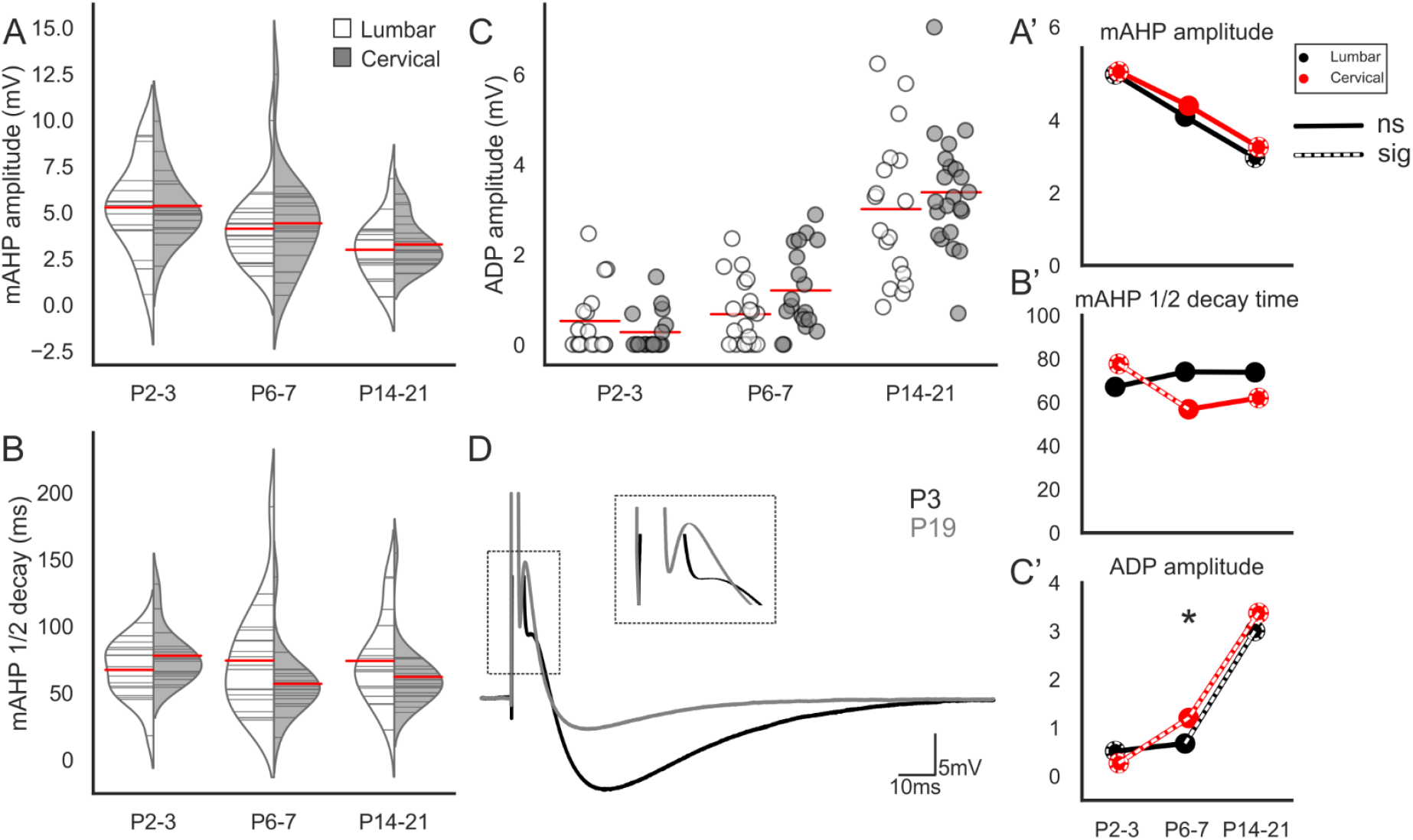
Postnatal development of after potentials. (A-B) Split violin plots illustrating the changes in mAHP amplitude and mAHP ½ decay time at each age for lumbar (white) and cervical (grey) motoneurons. The mAHP amplitude was measured from baseline (all cells held at −65 mV) to the most negative point of the mAHP. Half decay time was measured from the mAHP peak to baseline. (C) Strip-plots showing development of ADP amplitude with age. ADP amplitude was measured from the most negative point of the fAHP to the peak of the ADP-see inset with expanded ADPs. (D) Averaged trace (15-30 sweeps) of the action potential evoked with a 1ms square current pulse for a representative cell at P3 (black) and P19 (grey). (A-C’) Joint-plots illustrating the developmental profile of each measure. Over-laid white dashed lines represent statistically significant changes between age groups. Dashed circles represent statistically significant difference between P2-3 and P14-21.* represents a statistically significant difference between segments for a particular age group.

The mAHP 1/2 decay time did not change significantly during development of lumbar motoneurons (KW χ^2^= 0.43,p= 0.8055), but it did in cervical motoneurons (KW χ^2^= 14.1,p= 8.7e-03), where it reduced from 77.7 +/− 20.2ms at P2-3 to 57 +/− 24ms at P6-7 (p =8.4e-03). There was no difference between P6-7 and P14-21 (62 +/−24ms, p=0.338). As with the mAHP amplitude, there were no differences between lumbar and cervical motoneurons for decay time at any of the age groups (P2-3, p=0.432; P6-7, p=0.253; P14-21, p=0.202, Figure 3B, B’).

The ADP amplitude (measured from the trough of the fAHP to the peak of the subsequent depolarisation, if present) increased during development for lumbar and cervical motoneurons (Lumbar: KW χ^2^ = 32.0, p= 1.145e-07; Cervical: KW χ^2^ = 40.7, p= 1.461e-09, Figure 3C, C’, D). In fact, 41% of all motoneurons (regardless of segment) expressed an ADP at P2-3 (14/34), with this proportion increasing to 77% at P6-7 (30/39), and 100% of motoneurons at P14-21 (43/43). In the lumbar spinal cord, 47% (8/17) of motoneurons expressed an ADP at P2-3, 65% (13/20) at P6-7, and 100% (18) at P14-21. However, there was no difference between mean ADP amplitude between P2-3 (0.5 +/− 0.8mV) and P6-7 (0.7 +/− 0.7mV, p=0.311). Between P6-7 and P14-21, the ADP amplitude in lumbar motoneurons increased to 3.0 +/− 1.6mV (p= 8.2e-06). In the cervical spinal cord at P2-3, the proportion of motoneurons expressing an ADP was 41% (7/17) and the amplitude 0.3 +/− 0.4mV. This increased to 89% (17/19) and 1.2 +/− 0.9mV at P6-7 (p=2.1e-03) and to 100% (22) and 3.3 +/− 1.3mV between P6-7 and P14-21 (p= 1.2e-06). There was no difference in ADP amplitude between lumbar and cervical motoneurons at P2-3 (p=0.401) or P14-21(p=0.699), but it was greater in cervical Mns at P6-7 (p=0.044), consistent with the interpretation that maturation occurs earlier in the cervical spinal cord.

Given the importance of ADPs in generating high initial frequency firing frequencies and doublets, these data suggest development of ADPs enable motoneurons to fire at higher initial frequencies as they mature. Further, this increase occurred earlier in cervical motoneurons suggesting that development of high initial firing frequencies would be more mature compared to lumbar motoneurons at the same time points (see below).

### Postnatal development of motoneuron transition properties: sag potentials

Sag potentials mediated by Ih can also be considered to be transitional properties, as they contribute to motoneuron firing and are modulated by various neurotransmitters associated with input amplification (Ito & Oshima, 1965; Berger *et al.*, 1996). We assessed the slope of the sag potential, which is a reliable representative of the Ih conductance change (Figure 4A-D). In general, there seemed to be a reduction in the sag slope between P2-3 and P6-7 followed by an increase at P14-21. We saw no significant effect of age in lumbar motoneurons (KW χ^2^ = 5.7, p= 0.057) but there was a significant effect on cervical motoneurons (KW χ^2^= 15.1, p=5.0e-03). In cervical motoneurons there was a reduction between P2-3 (−6.7 +/− 8.0 mV/nA) and P6-7 (2.6 +/− 2.1 mV/nA, p= 0.042), and a significant increase between P6-7 and P14-21 (5.5 +/− 2.9mV/nA, p= 1.5e-03). There was no difference between segments at any age (P2-3, p=0.788, P6-7, p=0.231, P14-21, P=0.650.

**Figure 4.**
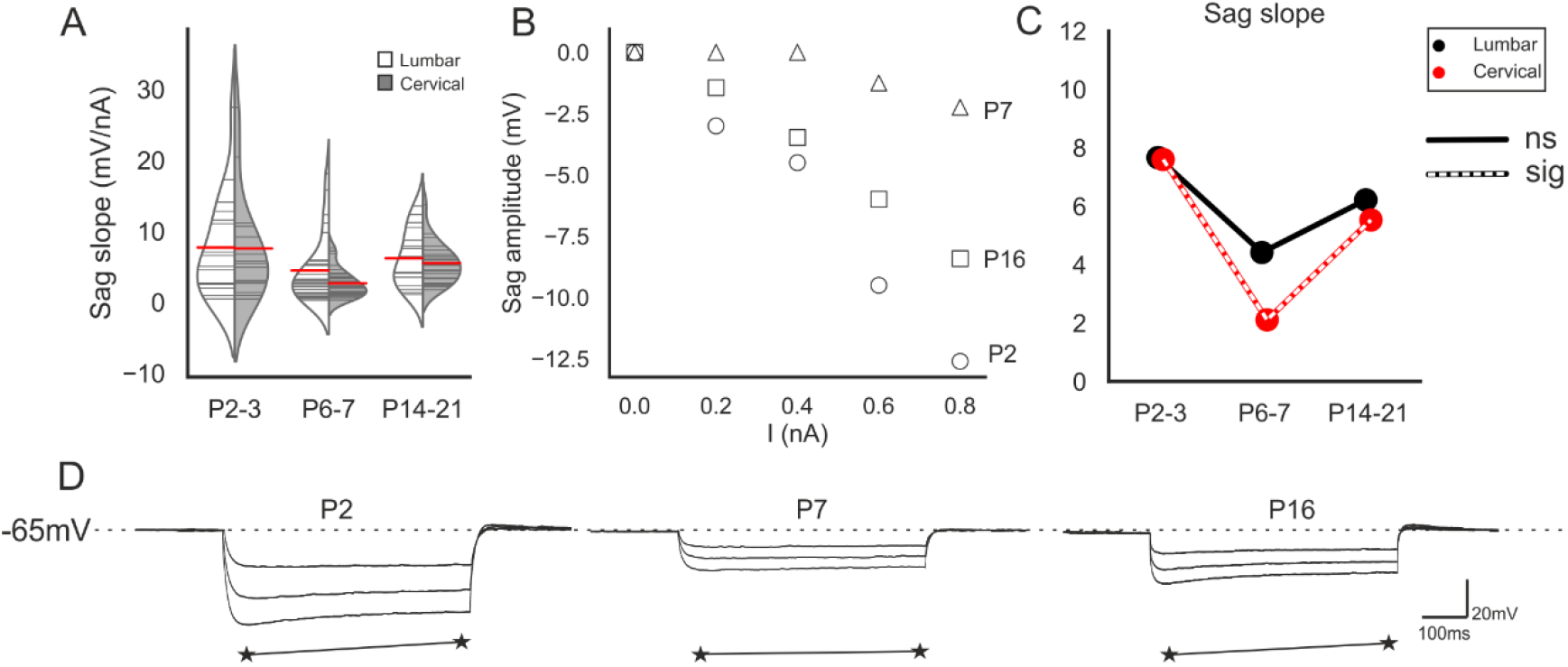
Postnatal development of sag potential. (A) Split violin plots illustrating the changes in sag slope with development. (B) Example plots of sag voltage vs current injection for 3 motoneurons illustrate the changes in sag conductance (inverses of slope). Joint-plots of sag slope means illustrating the developmental profile. Over-laid white dashed lines represent statistically significant changes between age groups. (D) Examples of voltage responses to −0.4, −0.7 and -InA current injections in representative motoneurons from each age group. The sag amplitude was measured as the difference between the negative voltage peak at the start of the 500ms pulse and the steady state at the end (stars).

### Postnatal development of motoneuron firing properties: Minimum current for repetitive firing (I_min_)

For function, motoneurons must fire repetitive trains of action potentials. The frequencies of firing are graded, with increased synaptic (or injected) current leading to higher rates of firing. By plotting the frequency of firing vs the current injected, various key measures can be quantified, including the minimum current needed for repetitive firing (I_min_), the maximum firing rate obtainable (F_max_) and the slopes for initial, steady state, and overall relationships.

Consistent with the observed increase in motoneuron WCC (i.e. size) and decrease in membrane resistance, there was a significant increase in I_min_ over postnatal development for both lumbar (KW χ^2^ = 15.7, p=3.8e-03) and cervical motoneurons (KW χ^2^ = 28.1, p= 8.1e-07, Figure 5A, A’). In lumbar motoneurons, there was no change in I_min_ between P2-3 (0.45 +/− 0.38nA) and P6-7 (0.52 +/−0.27nA, p=0.155), but it increased between P6-7 and P14-21 (1.50+/−1.11 nA, p= 0.003). In cervical motoneurons, I_min_ increased between P2-3 (0.37 +/− 0.20 nA) and P6-7 (0.81 +/− 0.45 nA, p=0.001), and further increased between P6-7 and P14-21 (1.29 +/−0.67 nA, p=0.012). The only difference observed between segments was the greater cervical motoneuron I_min_ at P6-7 (p=0.013), again consistent with these motoneurons maturing earlier than lumbar motoneurons (P2-3, p= 0.972; P14-21, p=0.720).

**Figure 5.**
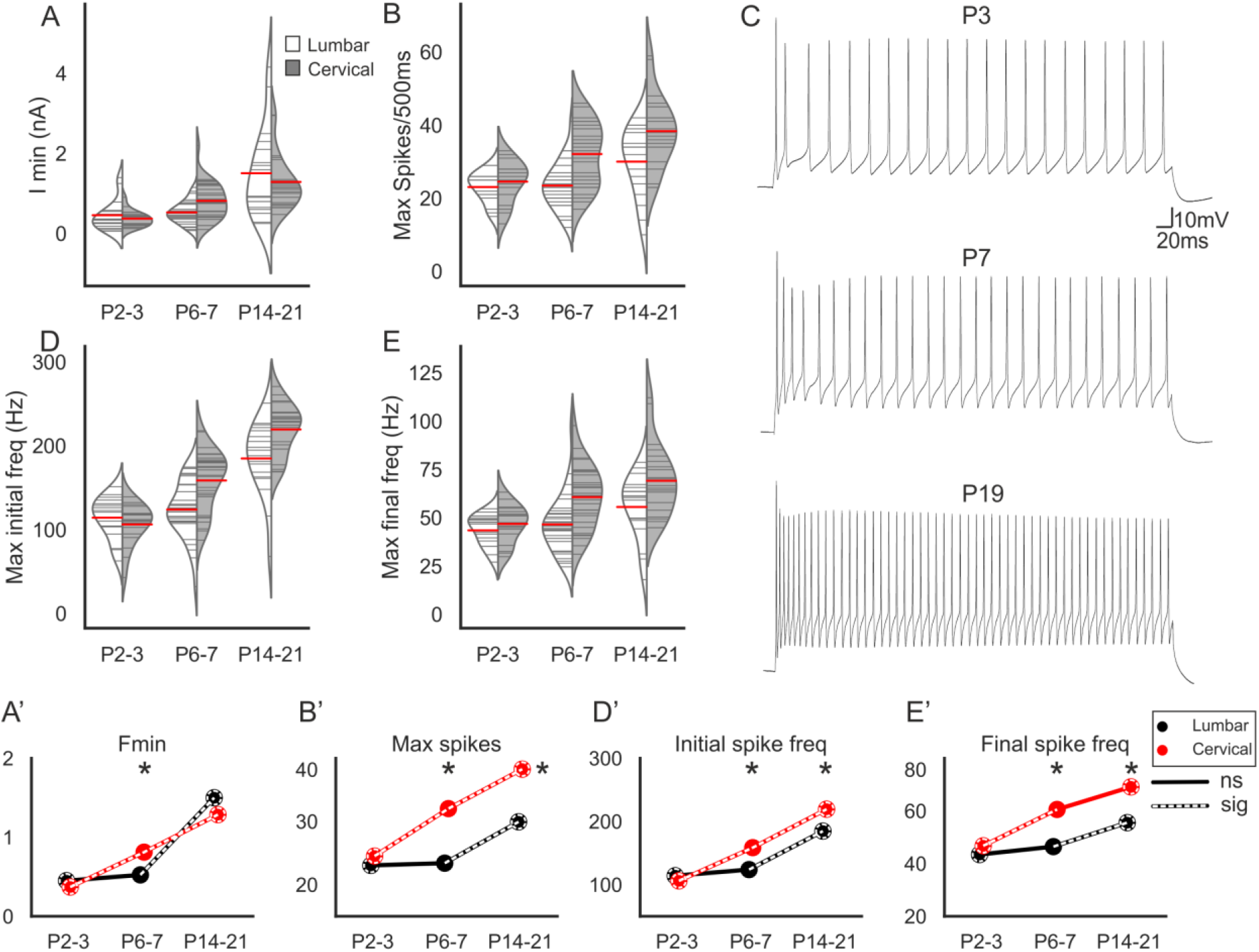
Postnatal development of repetitive tiring. (A-B) Violin plots showing the minimum current required for repetitive firing and maximum number of spikes in lumbar (white) and cervical (grey) motoneurons at each age group. (C) Representative traces of maximum firing of neurons from each age group illustrating the increased ability of motoneurons to produce high frequency trains of action potentials. Depolarising current pulses (500ms) of increasing magnitude were injected until the ccll reached its maximum firing frequency or depolarising block. (D) Max initial firing frequency was calculated from the first two action potentials at maximum firing rate. (E) The final firing frequency was calculated from the calculated from the last two action potentials at maximum firing rates. (A’-E’) Joint-plots of means illustrating the developmental profile for each measure. Over-laid white dashed lines represent statistically significant changes between age groups. Dashed circles represent statistically significant difference between P2-3 and PI4-21.* represents a statistically significant difference between segments for a particular age group.

### Postnatal development of motoneuron firing properties: Maximal spike output

In order to assess motoneuron firing capabilities throughout development, 500ms current pulses of increasing magnitude were injected into cervical and lumbar motoneuron until the maximum firing frequency (F_max_) was attained. In both the cervical and lumbar spinal cord, F_max_ increased with age (Figure 5B, B’, C). Specifically, the maximum number of spikes generated during a 500ms train increased with development for both lumbar (ANOVA, F=14.07, p=3.7e-03) and cervical motoneurons (ANOVA, F=23.64, p= 7.3e-06), as it did for maximum initial frequency (1^st^ 2 APs, Lumbar: ANOVA, F=48.11, p= 1.9e-09, Cervical: ANOVA, F=123.30, p= 2e-16, Figure 5D, D’) and maximum final frequency (final 2 APs, Lumbar: ANOVA, F= 9.24, p= 0.00337, Cervical: ANOVA, F=17.11, p= 1e-03, Figure 5E, E’)

In lumbar motoneurons, all measures of maximum firing increased between P6-7 and P14-21 (number of spikes per 500ms current pulse P6-7=23 +/− 6.5, P14-21= 30 +/− 9.4, p= 0.003; maximum initial freq: P6-7= 124 +/− 34 Hz,P14-21=185+/− 44 Hz, p= 9.7e-07; maximum final freq: P6-7=47 +/− 15 Hz, P14-21=56 +/− 17 Hz, p= 0.002). However, there was no difference between P2-3 and P6-7 for any of the measures (number of spikes per 500ms current pulse P2-3=23 +/− 4.2 Hz, p=0.902; maximum initial freq: P2-3=114 +/− 25 Hz, p= 0.357 or maximum final freq:P2-3=44 +/− 8.1 Hz, p= 0.734).

In cervical motoneurons, maximum initial frequency increased between P2-3 (106 +/− 25 Hz) and P6-7 (159 +/− 35 Hz, p=2.2e-05) and between P6-7 and P14-21 (219 +/− 30 Hz, p=3.9e-07). There was also an increase between P2-3 and P6-7 in both maximum spike number (P2-3= 25 +/− 5.8, P6-7= 32 +/− 8.7, p= 0.017) and final frequency (P2-3= 47 +/− 9.9, P6-7= 61 +/− 16, p= 0.007). However there was no difference between P6-7 and P14-21 for maximum final frequency (P14-21= 69 +/− 19 Hz, p= 0.186) and a small increase in maximum spike number (P14-21= 38 +/− 9.8, p=0.050).

A corrected pairwise comparison of cervical and lumbar motoneurons at P2-3 showed no differences for any of the maximum firing capabilities measured (number of spikes per 500ms current pulse, p= 0.355, maximum initial frequency, p=0.357; maximum final frequency, p=0.342). However, at both P6-7 and P14-21, mean cervical motoneuron values were greater than lumbar motoneurons for all measures (number of spikes per 500ms current pulse P6-7: p=0.001, P14-21: 0.019; maximum initial frequency P6-7: p=6e-03, P14-21: p=0.001; maximum final frequency P6-7: p=0.001, P14-21: p=0.033). Thus, repetitive firing parameters seem to mature earlier in cervical than in lumbar motoneurons.

Overall, motoneuron repetitive firing capacity increased with development. However, there seem to be differences in the profile of development between cervical and lumbar motoneurons. Specifically – and consistent with developmental profile of AP characteristics – cervical motoneurons increase their firing capabilities earlier in development compared to lumbar motoneurons and have higher maximum firing frequencies in the 3^rd^ postnatal week.

### Postnatal development of motoneuron firing properties: Excitability

Next, we assessed the changes in motoneuron excitability with development. This was done by measuring the slope of the linear portion of the relationships between the current injected and each of: number of spikes per 500ms pulse (spike slope); instantaneous frequency of the first two spikes (initial firing frequency); or instantaneous frequency of the last two spikes (final firing frequency or steady state, Figure 6 A-C). For cervical and lumbar motoneurons, age had a significant effect on all these measures (spike number f-I slope: Lumbar-KW χ^2^ = 30.0, p=3.0e-07, Cervical-KW χ^2^ = 32.1, p=1.1e-07; initial frequency gain: Lumbar-KW χ^2^ = 18.0, p=1.2e-03, Cervical-KW χ^2^ = 17.9, p=1.2e-04; final frequency gain: Lumbar-KW χ^2^ = 26. 7, p=1.6e-06, Cervical-KW χ^2^ = 36.7, p=1.1e-08).

**Figure 6.**
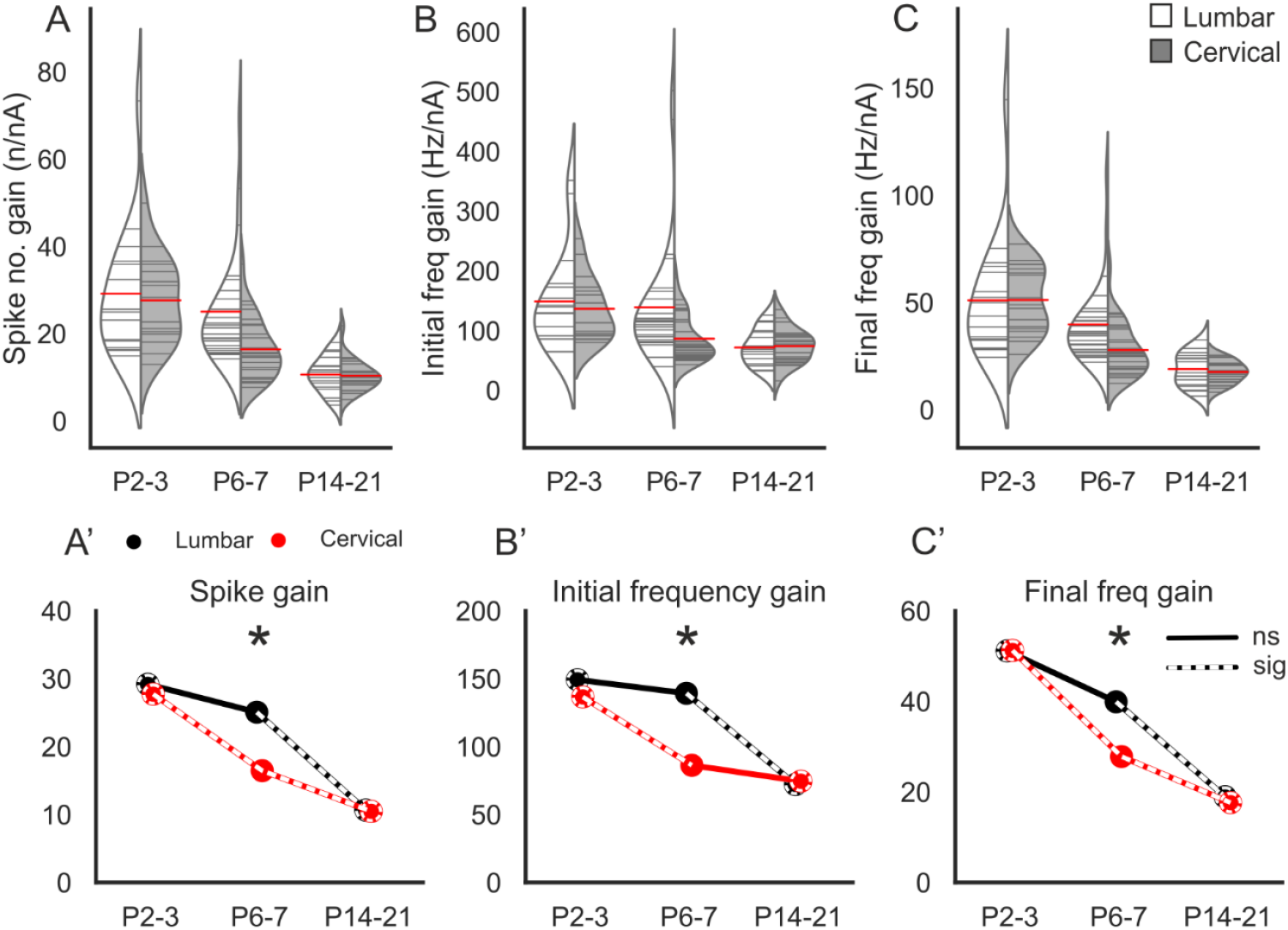
Postnatal development of cervical and lumbar motoneuron excitability. (A-C) Violin plots illustrating postnatal changes in the slope of the linear portion of an I/F plot. (A’-C’) Joint plots of means illustrating developmental profile of each measure. Over-laid white dashed lines represent statistically significant changes between age groups. Dashed circles represent statistically significant difference between P2-3 and P14-21. * represents a statistically significant difference between segments for a particular age group.

For lumbar motoneurons, there was no change in the spike f-I slope between P2-3 (29+/− 15 spikes/nA) and P6-7 (25 +/− 13 spikes/nA, p=0.305). However, there was a significant decrease between P6-7 and P14-21 (11 +/− 4.1 spikes/nA, p=7.1e-06). The spike slope in cervical motoneurons decreased between all age groups (P2-3= 30 +/− 10 spikes/nA vs P6-7= 17 +/− 6.8 spikes/nA, p=4.4e-03; P6-7 vs P14-21= 11 +/− 3.6 spikes/nA, p= 8e-03). There was no difference in spike slope between lumbar and cervical motoneurons at P2-3 (p=0.958) or P14-21 (p=0.359), however it was lower in cervical motoneurons at P6-7 (p=8e-03).

Initial firing frequency slope decreased in lumbar motoneurons between P6-7 (139 +/− 102 Hz/nA) and P14-21 (72 +/− 30 Hz/nA, p=5.4e-03), but there was no difference between P2-3 (149 +/− 84 Hz/nA) and P6-7 (p=0.504). In cervical motoneurons, there was a decrease in the slope of the initial frequency gain between P2-3 (137 +/− 53 Hz/nA) and P6-7 (86 +/− 37 Hz/nA, p=5.4e-03), but not between P6-7 and P14-21 (75 +/− 29 Hz/nA, p= 0.543).

Furthermore, as with the spike slope, initial frequency gain was lower in cervical than lumbar motoneurons at P6-7 (p= 0.003), but there was no difference at P2-3 (p=0.972) or P14-21(p= 0.751).

For the slope of the instantaneous frequency of the final 2 spikes, in lumbar motoneurons there was a decrease between P6-7 (40 +/− 19 Hz/nA) and P14-21(18 +/− 7.9 Hz/nA, p=2.8e-05), but not between P2-3 (51 +/− 29 Hz/nA) and P6-7 (p=0.136). In cervical motoneurons, the steady state slope decreased significantly between all age groups (P2-3= 51 +/− 16 Hz/nA vs P6-7= 28+/− 12 Hz/nA, p=1.2e-05; P6-7 vs P14-21=18 +/− 5.1 Hz/nA, p=7.8e-03). Cervical motoneurons had a smaller slope for steady state gain at P6-7 compared to lumbar motoneurons (p=0.002), but there was no difference between segments at P2-3(p=0.495) or P14-21(p=0.369).

In summary, for almost all measures of repetitive firing, it can be seen that at P2-3 lumbar and cervical motoneurons are the same, but there is a reduction of excitability for cervical motoneurons at P6-7 (Figure 6A’-C’). In lumbar motoneurons, this reduction is seen later, between P6-7 and 14-21, indicating that the reduction in excitability occurs at earlier postnatal periods in cervical motoneurons.

### Principal Component Analysis

Analysis of the individual variables above suggests that overall there is, as expected, a significant effect of age on development of motoneuron membrane properties and firing characteristics. Additionally, for many of the properties, we found a difference in the developmental profile between lumbar and cervical motoneurons, with cervical motoneurons maturing earlier than lumbar motoneurons. Due to the high dimensionality of the data, we used principal component analysis to map overall changes in motoneuron properties between ages and segments, reasoning that distinct clusters would indicate different stages of maturity. The first 2 principal components accounted for 52.9% of the variance (Supplementary figure 1A), with contributing variables in the 1^st^ component being mainly firing characteristics (Supplementary figure 1B) and 2^nd^ component mainly passive properties and some AP morphology characteristics (Supplementary figure 1C). The first component shifts progressively to more positive values with development, while there is minimal change in the 2^nd^ component. Figure 7 illustrates that 95% confidence ellipses for cervical motoneurons at P2-3 and lumbar motoneurons at P2-3 display a large degree of overlap, suggesting that they have similar firing characteristics and membrane properties. There is then a “jump” to the right in the properties of cervical motoneurons at P6-7, but this does not occur in lumbar motoneurons until P14-21: these two groups cluster at positive values along the 1^st^ principal component with their ellipses also showing overlap. There is then a further move to the right at P14-21 in the properties of cervical motoneurons. Thus, as shown with many of the individual parameter measurements, cervical and lumbar motoneurons are very much alike at birth, but cervical motoneurons mature earlier than those in the lumbar spinal cord.

**Figure 7.**
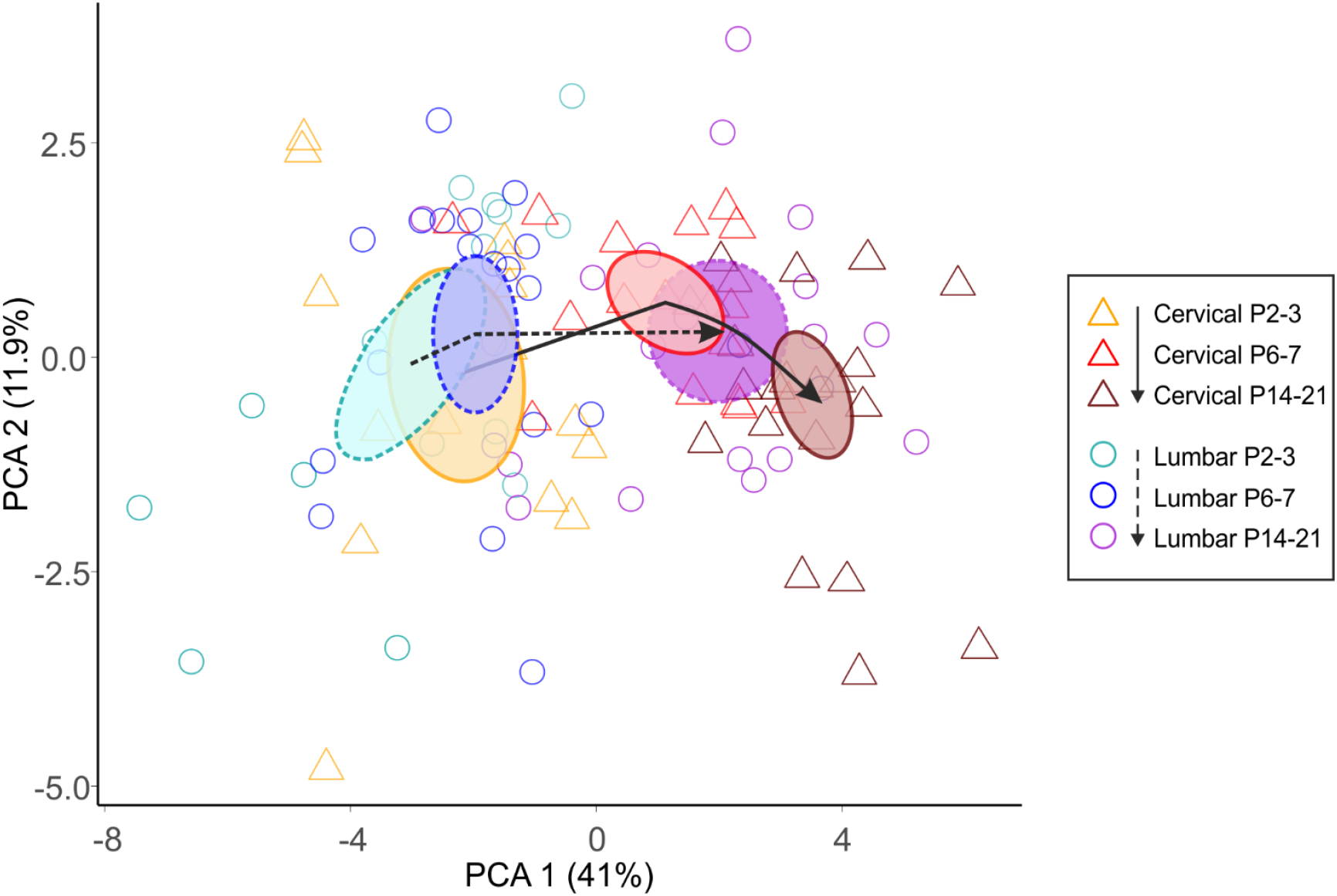
Principal component analysis of the development of motoneuron electrophysiological properties. The multidimentional dataset was reduced to principal components with the first 2 PCs accounting for 52.9 % of the variation, plotted. Ellipses were formed based on 95% confidence intervals for the group means for the I^st^ 2 components. Dashed ellipses represent the 3 age groups for lumbar motoneurons and the solid lines for cervical motoneurons. For individual points, triangle shapes represent cervical and circles represent lum bar motoneurons. Dashed and so lid arrows show the shift in the lumbar and cervical ellipsesrespectively during development.

## Discussion

Altricial mammals are born in an immature state, and their nervous systems and musculoskeletal systems must develop such that there is sufficient motor independence prior to the time of weaning. These two systems, which develop hand-in-hand, are connected by motoneurons and proprioceptive afferents. Here, we ask how the properties of motoneurons develop during this period, and ask whether the evidence supports cell autonomous or circuit factors as being the leading instigators for this development. To do this, we compare lumbar and cervical spinal motoneuron properties through this post-natal period, examining passive, transition, and repetitive firing properties from a stage where the animal is virtually immobile (P2-3) to the point when motor function matures, just prior to weaning (P14-21). We show that there is ongoing development of both cervical and lumbar motoneuron properties throughout this period and that despite the fact that cervical motoneurons are embryonically born prior to lumbar motoneurons (Nornes & Das, 1974), their properties at birth are similar. In the first 3 weeks of post-natal life, the properties of both sets mature, but those of cervical motoneurons develop earlier than those of lumbar motoneurons. This development correlates with the arrival of descending axons, which is known to be crucial to the maturation of spinal sensorimotor circuits. We therefore suggest maturation of these circuits contributes to development of motoneuron firing properties.

### The critical period: postnatal development of motoneuron properties continues into the 3^rd^ postnatal week

Weight bearing and fundamental aspects of motor control are established to a degree by P10-12, but it is clear that motor functional output in rodents does not mature until the 3^rd^ postnatal week (Altman & Sudarshan, 1975). Indeed, anatomical studies show that developmental organisation of membrane proteins and synaptic input to spinal motoneurons begin to reach maturity in the 3^rd^ week postnatally in rodents (Wilson *et al.*, 2004; Jean-Xavier *et al.*, 2017; Smith *et al.*, 2017). A profile of changes in passive properties and firing characteristics of motoneurons has been described for rodent hypoglossal and genioglossal motoneurons from birth to adulthood, but equivalent studies in the spinal cord have been limited to ages younger than P12 (Fulton & Walton, 1986; Nunez-Abades *et al.*, 1993; Berger *et al.*, 1996; Nakanishi & Whelan, 2010; Quinlan *et al.*, 2011). Due to the ready decline in motoneuronal viability in slices after P10, early neonatal spinal cord preparations have become the dominant tool to study motoneuron physiology, and therefore much of our knowledge is limited to this period in rodents and to the adult in cats (Kernell, 1999). Our recordings from spinal motoneurons in the cervical and lumbar spinal cord up to P21 confirm that maturation of electrophysiological properties continues into the 3^rd^ postnatal week, and demonstrates the importance of using older preparations to study neuronal properties.

Changes in motoneuron properties are dependent upon the expression and function of many membrane proteins. Although we did not study specific motoneuron ion channels, our results indicate that many channels undergo developmental changes in this critical period. During this time, WCC increases and input resistance is reduced, with a corresponding increase in I_min_. These changes are expected with growth of the motoneuron soma and dendritic tree, but also indicate that there is maturation of conductances that are active near resting potential (Fleshman *et al.*, 1981; Vinay *et al.*, 2000). Analysis of action potential morphology shows reduced half width due to increased rates of both depolarisation and repolarisation phases, suggesting that expression of Na^+^ (Barrett & Crill, 1980), Ca^2+^ (Hounsgaard & Mintz, 1988; Mynlieff & Beam, 1992; Viana *et al.*, 1993a), and K^+^ channels undergoes maturation (Viana *et al.*, 1993b). Thus, widespread changes to the motoneuron membrane occur during this period.

### Maturation of repetitive firing

A motoneuron’s raison d’être is to ensure that muscle fibres contract appropriately to produce the behaviour, and to do so the neuron must fire graded trains of action potentials. The maturation of spike properties, including increasing amplitude and reduced half width resulting from increasing rates of depolarisation and repolarisation, can facilitate faster firing (McCormick *et al.*, 1985). There is also an increase in the expression and amplitude of the ADP, a potential likely dependent upon high voltage activated Ca^2+^ channels (Granit *et al.*, 1963b; Kobayashi *et al.*, 1997; Vinay *et al.*, 2000). ADPs promote doublet firing in motoneurons, significantly increasing the rate and magnitude of muscle force generation (Parmiggiani & Stein, 1981; Sandercock & Heckman, 1997). The increase in ADP we report here may underlie the increase of maximum initial firing frequency across this developmental period. These changes in action potentials and ADPs thus likely underpin the changes in motoneuron repetitive firing capabilities.

The mAHP is also an important factor in regulating the frequency of firing, or interspike interval (Granit *et al.*, 1963a; Kernell & Monster, 1982; Bean, 2007; Deardorff *et al.*, 2013). This afterpotential is mediated by small conductance Ca^2+^ activated potassium channels (SK2 & SK3; Deardorff et al., 2013). We report a small developmental change in the mAHP amplitude in cervical but not lumbar motoneurons. Although greater changes in the mAHP might have been expected and have been reported in hypoglossal nuclei (Viana *et al.*, 1994; Berger *et al.*, 1996), our results are largely consistent with previous work in young spinal cord slices and brainstem slices throughout development (Carrascal *et al.*, 2005; Nakanishi & Whelan, 2010; Quinlan *et al.*, 2011).

The ability of motoneurons to fire trains of action potentials at high frequencies increased with development, while excitability, as measured by the gain of I/F plots, decreased. This reduction in I/F slope is likely related to the change in passive properties such as size of the soma and dendritic tree (Ulfhake & Cullheim, 1988; Ulfhake *et al.*, 1988), and active properties such as potassium channels associated with spike repolarisation (Gao & Ziskind-Conhaim, 1998; Martin-Caraballo & Greer, 2000). The combination of higher F_max_ and reduced I/F slope leads to a broader range of input currents to which the motoneuron responds, an increase in signal-to-noise ratio of the response, and an enhancement in the tunability motoneuron firing rates. It is intuitive that these properties continue developing throughout the 3^rd^ post-natal week and beyond, as the speed, force, and control of motor output continues to mature across this period (Altman & Sudarshan, 1975).

### Possibility of selection bias?

Could some of the differences in properties that we report result from selection bias? For example, SK2/3 expression and thus mAHP characteristics are different in different motoneuron types (i.e. those innervating fast vs slow twitch muscle fibres, which correspond to large vs small motoneurons (Deardorff *et al.*, 2013)). While in older preparations there is an inherent bias to record smaller motoneurons (because they survive; see Mitra and Brownstone (2012) where average input resistance was 123 MΩ), we were clearly not biased to smaller motoneurons: the mean input resistances we report in the oldest age group were approximately 30 MΩ. In fact, in this study, most motoneurons at P6-7 were under 50 MΩ and at P2-3 most were under 100 MΩ, both in the range of larger motoneurons. Furthermore, it seems that GFP expression decreases in Hb9::eGFP mice during this time period, persisting primarily in large motoneurons (and possibly even a specific subset of these). It is thus likely that we sampled the largest motoneurons in each age group. But there is also no doubt that motoneurons are growing during this period, and there is no way, at this point in time, to target neurons in younger mice that may be destined to grow up to be large motoneurons. That is, in order to reliably track the development of properties such as the mAHP, motoneuron types must be identified throughout development, which is a difficult proposition considering that motoneurons undergo significant diversification during postnatal development (Navarrete & Vrbová, 1993; Kanning *et al.*, 2010).

### Possible factors leading to advanced maturation of cervical vs lumbar motoneurons

Sequential rostro-caudal development of many components of the nervous system has been observed both pre- and postnatally. During embryonic development, spinal motoneurons are born first in the cervical spinal cord, and then sequentially in the more caudal segments (Nornes & Das, 1974). This could mean that delayed postnatal maturation of motoneuron firing characteristics in the lumbar cord may be set from the point of neurogenesis. We find this unlikely to be the case however, as we showed no difference between motoneuron properties in cervical vs lumbar segments at P2-3 for almost all the properties we analysed. However, by P6-7, cervical motoneurons had more mature properties than lumbar motoneurons. We therefore propose an alternative hypothesis suggesting that segmental differences in states of motor circuit maturity and activity patterns during postnatal development underlie the differences seen between cervical and lumbar motoneuron firing characteristics.

The delayed arrival and maturation (including myelination and synaptic refinement) of supraspinal descending systems is perhaps the most obvious difference between the two segments during postnatal development (Donatelle, 1977; Gianino *et al.*, 1999; Vinay *et al.*, 2005). And following arrival of these systems, their termination patterns as well as their synaptic effects on spinal motoneurons also mature - with cervical effects thus preceding lumbar effects (Commissiong, 1983; Tanaka *et al.*, 1992; Floeter & Lev-Tov, 1993; Brocard *et al.*, 1999b). This can certainly be seen in the development of postural control (Skoglund, 1960; Brocard *et al.*, 1999a), which occurs in a proximo-distal gradient corresponding to the more caudal location of motor nuclei innervating distal muscles (Nicolopoulos‐Stournaras & Iles, 1983). Furthermore, normal development of sensory afferents and spinal premotor circuits are dependent upon descending innervation (Chakrabarty & Martin, 2011b; Smith *et al.*, 2017), and these afferents themselves may affect motoneuron maturation (Woolley *et al.*, 1999). The importance of these systems on motoneuron properties has been seen when descending tracts are prevented from growing: there is disruption of the development of inhibitory systems such as Renshaw cells and GABA-pre circuits, and motoneuron hyperexcitability emerges (Martin, 2005; Chakrabarty *et al.*, 2009; Chakrabarty & Martin, 2011a; Chakrabarty & Martin, 2011b; Basaldella *et al.*, 2015; Smith *et al.*, 2017). We thus propose that rather than cell autonomous factors, the differences in the states of maturation of cervical vs lumbar motoneurons results from differences in the timing of local sensorimotor circuit development in the two regions, which is affected by innervation by descending systems.

### Conclusion

We show here that between birth and weaning, cervical motoneuron properties mature earlier than those of lumbar motoneurons. We suggest that this maturation results from the development of circuits that engage these motoneurons. It has been shown that the properties of many different neuronal types mature in this postnatal period, including thalamocortical neurons (Warren & Jones, 1997), somatosensory cortical inhibitory neurons (Kinnischtzke *et al.*, 2012), medial prefrontal cortex pyramidal neurons (Favuzzi *et al.*, 2019), and auditory cortex pyramidal neurons (Oswald & Reyes, 2008), amongst others. While of course cell autonomous factors such as transcription factor expression respond to environmental cues such as morphogens to govern cell fate, including the fate to become a motoneuron (Jessell, 2000b; Dasen *et al.*, 2008), the degree to which these molecules set a maturating course is not clear (Harb *et al.*, 2016). In fact, basic motoneuron properties can develop in motoneurons derived in a dish from embryonic stem cells (Miles *et al.*, 2004), demonstrating the strength of intrinsic programmes in determining electrophysiological properties. From our study of geographically separated motoneurons, however, we argue that circuit factors play an important role in physiological maturation. It seems likely that the combination of cell-autonomous factors and circuit development are required to ultimately produce a mature, functional neuron.

## Methods

### Ethical approval

Experiments were approved by the University College London Animal Welfare and Ethical Review Body and performed under a license granted by the Home Office Animals (Scientific Procedures) Act, 1986.

### Animals

All experiments were carried out on Hb9::eGFP mice of both sexes aged P2-P21. Male Hb9::eGFP (B6.Cg-Tg(Hlxb9-GFP)1Tmj/J) mice were acquired initially from Tom Jessell’s lab and have been bred in the Brownstone lab since the early 2000s. This strain is available from JAX (stock no. 005029). We breed them with C57/Bl6 wild type females acquired from Charles River Laboratories, Inc (strain code:027). The date of birth was called P0 and recordings were made at 3 different age ranges (P2-3, P6-7 and P14-21). A total 54 animals were used in the study (lumbar P2-3, n=7; cervical P2-3, n=6; lumbar P6-7, n=11; cervical P6-7, n=11; lumbar P14-21, n= 8, cervical P14-21, n=11). Average age of mice from which P14-21 lumbar motoneurons were sampled was 17.1 ± 2.0 days and 16.2 ± 1.7 days for cervical motoneurons. Motoneurons from lumbar or cervical segments were always sampled from separate mice.

### Spinal cord isolation

#### Lumbar spinal cords

Animals were deeply anaesthetised by intraperitoneal injection of a ketamine (100mg/kg) and xylazine (20mg/kg) mixture. Upon loss of paw withdrawal, mice were then decapitated and the vertebral column was quickly isolated and pinned down (ventral-side-up) in a dissecting dish containing ice cold (0-4°C) normal artificial cerebrospinal fluid (nACSF) saturated with 95% carbogen. The composition of the nACSF was as follows (in mM): 113 NaCl, 3 KCL, 25 NaHCO_3_, 1NaH_2_PO_4_, 2 CaCl, 2 MgCl_2_ and 11 D-glucose, pH 7.4 (Mitra & Brownstone, 2012). The vertebral bodies were removed to reveal the spinal cord, the roots were cut and dura matter removed. The spinal cord was isolated from rostral thoracic to caudal sacral segments, placed upon tissue paper to soak up excess liquid and then the dorsal side glued (3M Vetbond^TM/MC^, No.1469SB) to a pre-cut block of agarose mounted on a cutting chuck.

#### Cervical spinal cords

For cervical slices (also under deep anaesthesia), a craniotomy was done to expose and remove the cerebellum, and the brainstem was transected at the level of the obex. All nervous tissue rostral to the transection was immediately removed. This process was critical for preserving cervical spinal tissue. The vertebral column was then transferred to a dissecting dish (as with lumbar preparation), pinned dorsal-side-up, and a laminectomy performed to a mid-thoracic level. Roots were cut, the dura removed and cervical cord glued to the chuck in the same fashion as the lumbar cord.

### Slice preparation

The cord was transferred to the vibrating microtome (Model 7000smz-2, Campden Instruments Ltd.) chamber containing ice cold slicing solution made up of the following (in mM): 130 potassium gluconate, 15 KCL, 0.05 EGTA, 20 HEPES, 25 glucose, 3 kynurenic acid, pH 7.4 (Dugué *et al.*, 2005; Bhumbra & Beato, 2018). Slices were made at 350μm, transferred to a recovery chamber containing nACSF (32°C) for 30 minutes and then left to equilibrate to room temperature for at least 30 minutes before recording (1 hour total post slice recovery).

### Electrophysiology

Motoneuron recordings were made with a MultiClamp 700A amplifier (Axon Instruments, Inc), low pass filtered at 10kHz and digitized at 25kHz using a CED Power3 1401 and Signal software (Cambridge Electronic Design Ltd, Cambridge, UK). Patch pipette electrodes were pulled with a horizontal puller (P-97 Flaming/Brown Micropipette Puller; Sutter Instrument) to a resistance of 1-5 MΩ and filled with the internal recording solution made up of (in mM): 131 K-methanesulfonate, 6 NaCl, 0.1 CaCl_2,_ 1.1 EGTA, 10 HEPES, 0.3 MgCl_2_, 3 ATP-Mg, 0.5 GTP-Na, 2.5 L glutathionine, 5 phosphocreatine, pH 7.25 adjusted with KOH, osmolarity 290-300 mOsm.

Large eGFP positive neurons in the ventral lumbar and cervical spinal cord were identified as motoneurons (Wilson *et al.*, 2005). Motoneurons were patched at room temperature (approx 21°C) using infrared-differential interference contrast (IR-DIC) optics on a DMLFSA microscope (Leica DMLFSA; Leica Microsystems). All lumbar motoneurons were selected from the lateral motor nuclei of segments L1-6 and all cervical cells were selected from the lateral motor nuclei of slices made from C4-8. It is therefore likely that the majority (if not all) of cells included in our study were limb innervating motoneurons (Watson *et al.*, 2009; Mohan *et al.*, 2014; Mohan *et al.*, 2015).

### Data analysis

All data were captured and analysed with CED Signal software. Motoneurons were only analysed if they had a resting membrane potential more hyperpolarised than −60 mV that did not deviate by more than 5mV during recording.

All experiments were performed in current clamp mode. The bridge was balanced and capacitance neutralised prior to commencing recording. Motoneurons were injected with a small negative rectangular pulse (500ms duration) and the voltage response of 15-30 traces was averaged to measure input resistance and whole cell capacitance. Resistance was measured as the peak voltage change to the injected current and τ calculated from an exponential curve fitted to the response (automated in Signal). Whole cell capacitance was calculated using resistance and τ values and cross-checked against the values automatically recorded by the software during the experiment. I_min_ was defined as the minimum amount of current needed to evoke >= 2 action potentials. This was tested with 500ms duration, rectangular current pulses of increasing magnitude (0.03nA steps) starting at −0.3nA. Sag potentials were recorded by injecting 0.03nA hyperpolarising current steps (500ms) from 0 to −1nA. The sag amplitude was measured as the difference between the peak of the negative voltage deflection at the start of the 500ms pulse and the steady state at the end. Motoneuron I/F graphs were generated by injecting depolarising current steps increasing from 0nA until maximum firing was observed. The excitability of the cell (gain) was determined by measuring the slope of the linear portion of I/F plots for spike number, initial frequency (first 2 spikes instantaneous frequency), and final frequency (steady state, final 2 spikes instantaneous frequency).

Action potential half widths (AP HWs), spike amplitude, maximum rate of depolarisation/repolarisation and fast AHPs (fAHP) were measured from 15-30 averaged single APs evoked with a 20ms rectangular current pulse. The AP HW was calculated as the time between the 50% rise and 50% fall in amplitude of the AP. Spike height was measured as the voltage difference between the threshold (voltage at maximum positive value of the 2^nd^ derivative of membrane potential of AP) and the peak of the AP. The fAHP was measured as the difference between the voltage baseline and the most negative point on the 1^st^ trough of the AP. Phase-plane plots of the single APs were generated to calculate maximum rates of depolarisation and repolarisation. After-potential measurements (mAHP, mAHP ½ decay time and ADP amplitude) were taken from single APs evoked with a 1ms duration current pulse. The mAHP amplitude was calculated from baseline (held at −65 mV) to the most negative point on the trough. The mAHP ½ decay time is calculated as half the time taken (ms) from most negative point of the mAHP to baseline. After depolarisation (ADP) amplitude was measured from the fAHP peak to the most positive value before the start of the mAHP repolarisation. All cells were held at −65mV for single action potential experiments.

### Statistics

Data from all cells was initially exported to a Microsoft Excel file. All subsequent data processing and analysis was done using either RStudio or Python running through the Jupyter notebooks environment. Given the diversity of motoneurons in the spinal cord and in line with the 3 Rs for animal research, measures from individual cells were treated as the experimental unit (N).

The number of cells analysed for each measure can be found in supplementary tables 1–3. To assess effects of age and spinal segment, data was initially tested for normality (Shapiro wilks) and then either one way ANOVA or Kruskal-Wallis rank sum tests were performed depending on the outcome of the normality test. We first assessed if there was an effect of age on electrical properties of cervical and lumbar motoneurons separately. Unless there was no effect of age on either segment, we proceeded to compare the 6 groups with each other (2 regions, each at 3 ages) using either 1 way ANOVA or Kruskal-Wallis rank sum tests with pairwise comparisons. Pairwise comparisons were also performed based on normality (T-tests or Wilcoxon rank sum test) and p values were adjusted using the Bonferonni-Holm method. All data are reported as means and standard deviations. Graphs were produced using Seaborn and Python open source software. Split violin plots with individual observations (grey lines) and means (red) were used to show and compare the distribution of the data.

Schematic line plots are based on the means of the data and were included to give a clear illustration for comparison of the developmental profile between cervical and lumbar segments. Final figures were produced using CorelDraw Home & Student X8 software.

### PCA Analysis

Principal component analysis was performed in R studio using packages prcomp and Factoextra. First, the variables were scaled in order to standardise the data before generating the loadings (Lê *et al.*, 2008). Individual values were projected onto the 2D subspace and 95% CI ellipses were generated using the Factoextra package. The 95% CI was based on the the values from the first 2 components.

## Acknowledgements

We would like to thank Prof Marco Beato for valuable comments on a draft of the manuscript, and Nadine Simons-Weidenmaier, who helped to set up experiments and manage our mouse colonies. We are also grateful to Rafaela Fernandez De La Fuente for her help with managing the mouse colony. This work was supported by Wellcome (110193), and RB is supported by Brain Research UK.

## Contributions

CCS conceptualised and designed the study, performed the experiments and analysed the data; CCS and RMB interperted the data and wrote the paper.

## Supplementary Data

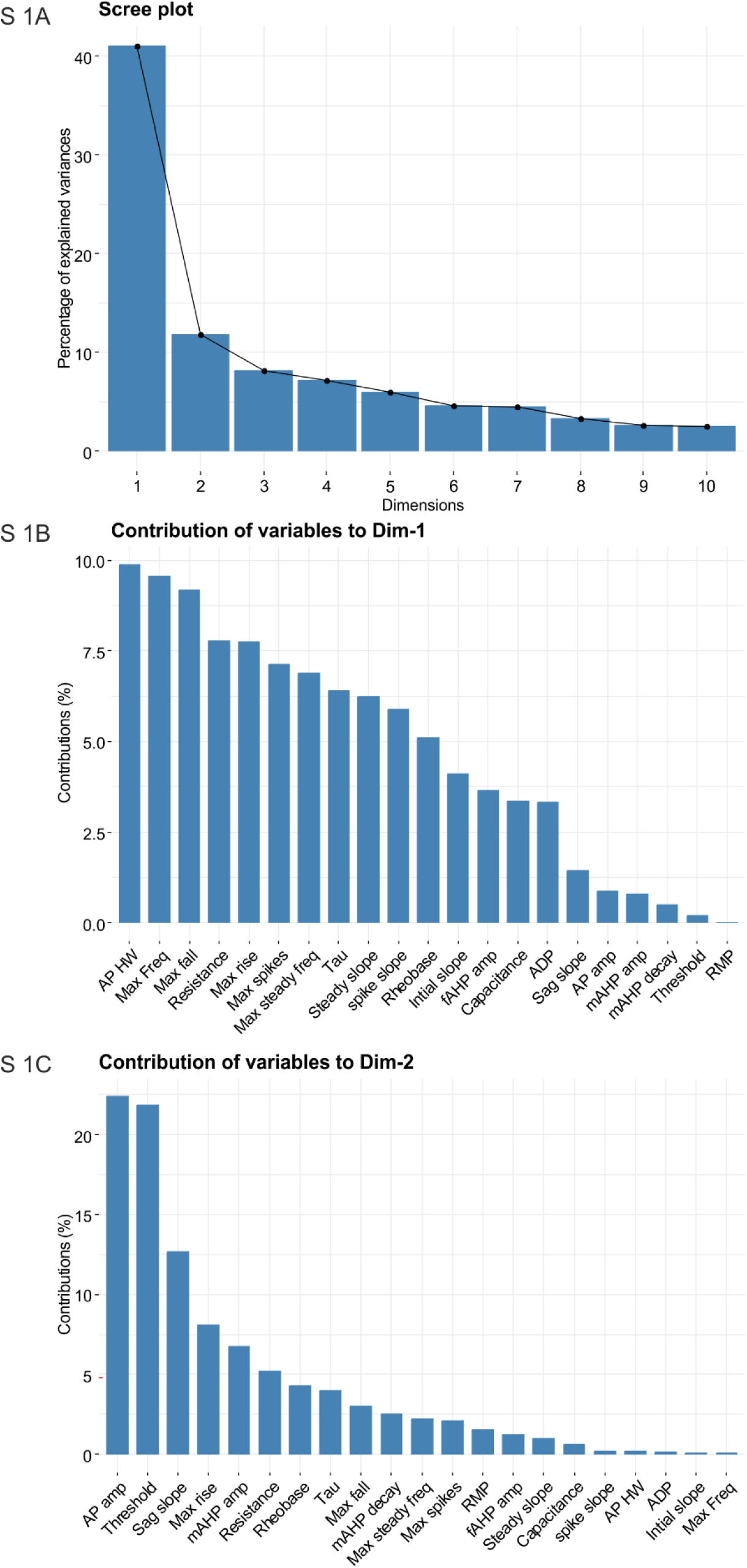

**Table. S1.** Postnatal development of passive membrane properties.

| Table S1. Postnatal development of passive membrane properties |  |  |  |  |  |  | 970 |
| --- | --- | --- | --- | --- | --- | --- | --- |
| Variable | Segment | P2-3 (n) | SD | P6-7 (n) | SD | P14-21 (n) | SD<sub>971</sub> |
| RMP (mV) | L | -65 | 4.4 | -65 | 3.2 | -67 | 2.7<sub>972</sub> |
|  | C | -67 | 3.1 | -65 | 2.8 | -66 | 2.4 |
| WCC (pF) | L | 223 | 71 | 310 | 140 | 405 | 217<sub>973</sub> |
|  | C | 212 | 59 | 353 | 136 | 340 | 148 |
| Res (MΩ) | L | 79 | 47 | 52 | 32.0 | 31 | 23<sub>974</sub> |
|  | C | 74 | 31 | 43 | 34 | 25.5 | 7.5 |
| Tau (ms) | L | 16.5 | 6.7 | 13.5 | 4.9 | 9.6 | 3.9 |
|  | C | 14.8 | 4.7 | 12.9 | 5.9 | 8.2 | 3.3 |
| (n) | L | 17 |  | 30 |  | 18 |  |
|  | C | 17 |  | 30 |  | 22 |  |

**Table. S2.** Postnatal development of transition membrane properties.

| Table S2. Postnatal development of transition membrane properties |  |  |  |  |  |  |  |
| --- | --- | --- | --- | --- | --- | --- | --- |
| Variable | Segment | P2-3 (n) | SD | P6-7 (n) | SD | P14-21 (n) | SD |
| Sag Slope (mV/nA) | L | 7.7 | 7.3 | 4.4 | 4.8 | 6. | 4.0 |
|  | C | 6.7 | 8.0 | 2.1 | 2.6 | 5.5 | 2.9 |
| Early action potential characteristics |  |  |  |  |  |  |  |
| AP HW (ms) | L | 1.2 | 0.2 | 1.0 | 0.2 | 0.8 | 0.2 |
|  | C | 1.1 | 0.2 | 0.8 | 0.2 | 0.6 | 0.1 |
| Depol rate (mV/s) | L | 152 | 33 | 166 | 34 | 212 | 53 |
|  | C | 150 | 51 | 192 | 44 | 255 | 46 |
| Repol rate (mV/s) | L | 65.9 | 13.7 | 80.7 | 15.3 | 115 | 34.1 |
|  | C | 77 | 22 | 100 | 28 | 149 | 35 |
| AP amp (mV) | L | 69 | 7.8 | 68 | 6.1 | 70 | 8.5 |
|  | C | 63 | 9.2 | 69 | 7.4 | 73 | 7.7 |
| Threshold (mV) | L | -36 | 9.5 | -37 | 7.2 | -35 | 8.2 |
|  | C | -37 | 8 | -34 | 8.3 | -35 | 6.0 |
| fAHP (mV) | L | -8.6 | 7.5 | -9.9 | 3.5 | -16 | 3.6 |
|  | C | -11 | 4.1 | -14 | 4.8 | -19 | 5.5 |
| (n) | L | 17 |  | 30 |  | 18 |  |
|  | C | 17 |  | 30 |  | 22 |  |
| Action potential late after potentials |  |  |  |  |  |  |  |
| mAHP amp (mV) | L | 5.3 | 2.4 | 4.1 | 1.9 | 3.0 | 1.6 |
|  | C | 5.4 | 1.9 | 4.4 | 2.6 | 3.3 | 1.3 |
| mAHP decay (ms) | L | 67 | 22 | 74 | 39 | 74 | 31 |
|  | C | 78 | 20 | 57 | 24 | 62 | 24 |
| ADP amp (mV) | L | 0.5 | 0.8 | 0.7 | 0.7 | 3.0 | 1.6 |
|  | C | 0.3 | 0.5 | 1.2 | 0.9 | 3.4 | 1.3 |
| (n) | L | 17 |  | 20 |  | 18 |  |
|  | C | 17 |  | 19 |  | 22 |  |

**Table. S3.**
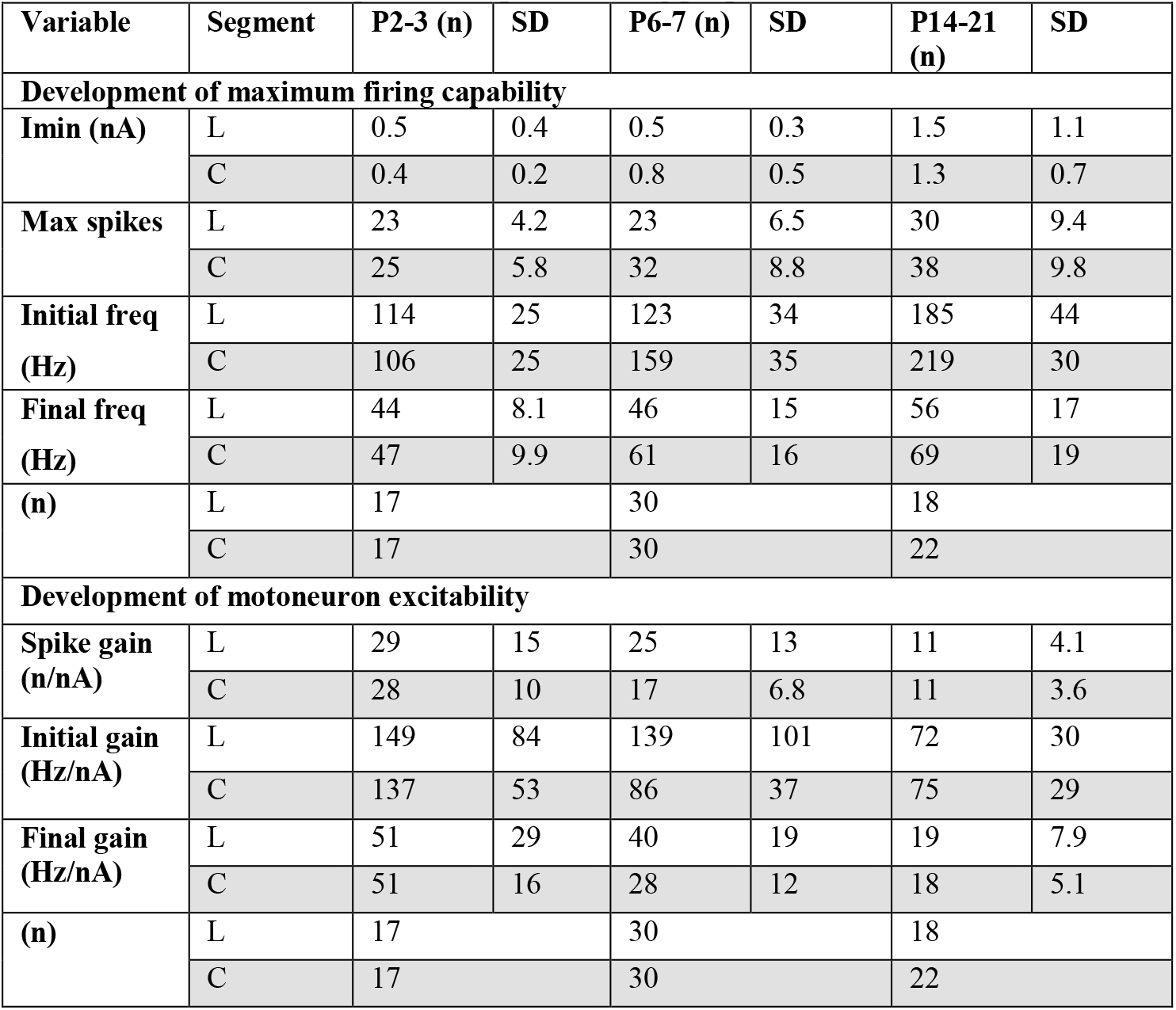
Postnatal development of repetitive firing properties.

| Table S3. Postnatal development of repetitive firing properties |  |  |  |  |  |  |  |
| --- | --- | --- | --- | --- | --- | --- | --- |
| Variable | Segment | P2-3 (n) | SD | P6-7 (n) | SD | P14-21 (n) | SD |
| **Development of maximum firing capability** |  |  |  |  |  |  |  |
| Imin (nA) | L | 0.5 | 0.4 | 0.5 | 0.3 | 1.5 | 1.1 |
|  | C | 0.4 | 0.2 | 0.8 | 0.5 | 1.3 | 0.7 |
| Max spikes | L | 23 | 4.2 | 23 | 6.5 | 30 | 9.4 |
|  | C | 25 | 5.8 | 32 | 8.8 | 38 | 9.8 |
| Initial freq (Hz) | L | 114 | 25 | 123 | 34 | 185 | 44 |
|  | C | 106 | 25 | 159 | 35 | 219 | 30 |
| Final freq (Hz) | L | 44 | 8.1 | 46 | 15 | 56 | 17 |
|  | C | 47 | 9.9 | 61 | 16 | 69 | 19 |
| (n) | L | 17 |  | 30 |  | 18 |  |
|  | C | 17 |  | 30 |  | 22 |  |
| **Development of motoneuron excitability** |  |  |  |  |  |  |  |
| Spike gain (n/nA) | L | 29 | 15 | 25 | 13 | 11 | 4.1 |
|  | C | 28 | 10 | 17 | 6.8 | 11 | 3.6 |
| Initial gain (Hz/nA) | L | 149 | 84 | 139 | 101 | 72 | 30 |
|  | C | 137 | 53 | 86 | 37 | 75 | 29 |
| Final gain (Hz/nA) | L | 51 | 29 | 40 | 19 | 19 | 7.9 |
|  | C | 51 | 16 | 28 | 12 | 18 | 5.1 |
| (n) | L | 17 |  | 30 |  | 18 |  |
|  | C | 17 |  | 30 |  | 22 |  |

